# Massive surface membrane expansion without involvement of classical exocytic mechanisms

**DOI:** 10.1101/249284

**Authors:** Michael Fine, Christopher Bricogne, Donald W. Hilgemann

**Affiliations:** University of Texas Southwestern Medical Center, Department of Physiology, Dallas, Texas, USA; UCL Cancer Institute, University College London, Gower St, London, UK.

## Abstract

Activation of TMEM16F, a Ca^2+^ -dependent ion channel and lipid scramblase, causes massive surface membrane expansion in multiple cell types by unresolved mechanisms. We describe here that membrane expansion reflects opening of deeply invaginating surface membrane compartments when anionic phospholipids are lost from the cytoplasmic membrane leaflet. Compartments that open contain vesicle-associated membrane proteins (VAMPs) and can open with as little as one micromolar free Ca_i_^2+^. Cationic peptides that sequester anionic phospholipids open the compartments from the cytoplasmic side without Ca^2+^. Monovalent cations facilitate membrane expansion via coupled permeation with anionic phospholipids through TMEM16F. When monovalent cation concentrations are reduced, membrane expansion can be reversed by changing ion gradients and membrane voltage. Depolarization closes the compartments by generating inward cation gradients through TMEM16F that promote influx of anionic phospholipids. In summary, TMEM16F-mediated membrane expansion likely does not reflect exocytosis but rather the relaxation of constrictions that close surface membrane invaginations.

**Summary:** The surface membrane of diverse cell types can be remodeled by opening and closing surface invaginations that are held shut by proteins that bind negatively charged lipids and constrict the orifices of these compartments.

## Introduction

Plasma membrane (PM) dynamics in eukaryotic cells reflect to a large extent the function of exocytosis and endocytosis. In brief, the PM is continuously modified by discrete membrane fusion and excision events that transfer chemicals to and from the extracellular space and modify the protein content of the PM (Sudhof and Rothman, 2009). Beyond the transport of chemicals and membrane proteins, PM expansion can become essential for cell growth (Tojima et al., 2014), to allow cell swelling (Groulx et al., 2006), to repair PM wounds (Togo et al., 1999; Cooper and McNeil, 2015; Corrotte et al., 2015; Moe et al., 2015), to relax membrane tension caused by chronic stretch, e.g. bladder extension (Lewis and de Moura, 1984), and to execute multiple steps of fertilization in mammals (Gadella and Evans, 2011).

Both electrical and optical methods have revealed that diverse cell types increase their PM area substantially during large elevations of cytoplasmic Ca^2+^ (Coorssen et al., 1996; Ninomiya et al., 1996; Kasai et al., 1999; Togo et al., 2003; Kreft et al., 2004; Bretscher, 2008; Cohen et al., 2008; Yaradanakul et al., 2008). PM expansion can double PM area within a few seconds in some cell types (Wang and Hilgemann, 2008; Yaradanakul et al., 2008; Bricogne, XXXX). To what extent the different PM responses occur by a single molecular mechanism remains unclear. Ca^2+^-activated PM expansion in BHK fibroblasts is not inhibited by tetanus toxins and does not appear to use synaptotagmins as Ca^2+^ sensors (Wang and Hilgemann, 2008). In contrast to the usual function of synaptotagmins (Sudhof, 2012), PM expansion in fibroblasts has a nearly linear dependence on cytoplasmic Ca^2+^ (Yaradanakul et al., 2008). As described in a companion article (Bricogne, XXXX), PM expansion in Jurkat T-cells and HEK293 cells can be initiated by the transmembrane protein, TMEM16F (Bricogne, XXXX). When activated by Ca^2+^, TMEM16F evidently functions simultaneously as a nonselective cation channel and as a phospholipid scramblase (Yang et al., 2012; Suzuki et al., 2013; Fujii et al., 2015; Scudieri et al., 2015). We analyze here how these functions mediate PM expansion.

To facilitate the presentation of Results, we illustrate our working hypotheses in Fig. 1. As described in the companion article (Bricogne, XXXX), the rate of PM expansion in Jurkat T-cells, monitored via membrane capacitance (C_m_), mirrors the rise and fall of TMEM16F channel openings, monitored as membrane conductance (G_m_). As shown in Fig. 1A, the integral of the G_m_ signal (∫G_m_) can be scaled to match closely the rising C_m_ signal. Our explanation is that TMEM16F activity limits PM expansion, and this is supported by the phenotype of HEK293 cells that overexpress TMEM16F. Wild-type (WT) HEK293 cells have a low TMEM16F conductance and expand their PMs to only to a small extent during Ca^2+^ elevations, but overexpression of TMEM16F markedly enhances the PM expansion responses ((Bricogne, XXXX),Fig.5D).

**Figure 1.**
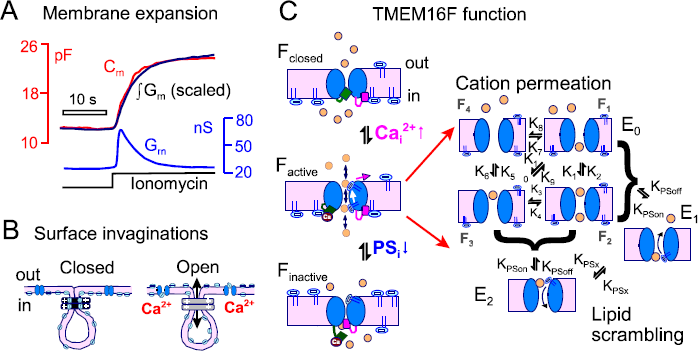
Working hypotheses to account for TMEM16F-mediated PM expansion. **(A)** Routine recording of membrane capacitance (C_m_) and conductance (G_m_) changes during a cytoplasmic Ca^2+^ elevation induced by ionomycin (5 µM) with 2 mM extracellular Ca^2+^ in WT Jurkat T-cells. Same conditions as in Fig. 2A-D and Fig. 9. As in >20 similar observations, the wave-form of C_m_ is superimposable with the integral of the G_m_ signal. **(B)** Membrane compartments that expand the cell surface extend deeply into cells and are closed by proteins that bind anionic phospholipids, depicted as light blue ovals. Loss of anionic phospholipids from the cytoplasmic leaflet relaxes the constrictions and opens the compartments to the extracellular space. **(C)** TMEM16F channels (blue) are closed (F_Closed_) and do not conduct ions or lipids in the absence of cytoplasmic Ca^2+^. When cytoplasmic Ca^2+^ rises, TMEM16F binds Ca^2+^ and opens with a nearly linear dependence on Ca^2+^ concentration (F_active_). Anionic phospholipids can enter the active TMEM16F channels (see expanded diagram) only when a monovalent cation (orange circle) is bound within the channel near the relevant pore opening. Anionic phospholipids permeate the channel together with monovalent cations in an electroneutral fashion. Accordingly, cation gradients can influence the movement of anionic phospholipids. In the presence of ion gradients, lipid scrambling can become voltage dependent in dependence on occupation of the channel by cations. TMEM16F channels enter an inactive state (F_inactive_) when PS is lost from cytoplasmic binding sites. Double inactive states without Ca^2+^ or PS bound are not depicted. The rate constants indicated in Fig. 1C, the fractional TMEM16F occupancy by ions (F_1_ to F_4_), and the fractional occupancies by anionic phospholipids (E_0_, E_1_ and E_2_) are employed in simulations presented in Results and described in Materials and Methods.

Fig. 1B illustrates how TMEM16F may mediate PM expansion. As documented in Results, the membrane compartments that expand the cell surface can extend quite deeply into cells, and they appear to be constricted in regions where they converge with the outer PM. Both openings and subsequent closings of these compartments are shown to be reversible. This requires that the compartments open without collapse of their structures, similar to proposals for ‘kiss-and-run’ exocytosis (Alabi and Tsien, 2013; Oleinick et al., 2017; Wen et al., 2017) and dense core exocytosis in chromaffin cells (Fulop et al., 2005). To explain that the compartments can close reversibly, we are forced to consider the hypothesis that the compartments are contiguous with the PM both before they open and after they close. As illustrated, this can be explained if the compartments are constricted by membrane proteins that bind anionic phospholipids, thereby isolating them electrically from the extracellular space. Loss of anionic lipids during TMEM16F activity would then release the constrictions and open the compartments.

To explain TMEM16F function, as described in Results, we propose that TMEM16F exists in three activity states; closed, open and inactive (Fig. 1C). In the absence of cytoplasmic Ca^2+^, TMEM16F channels are closed and have no scramblase activity. Upon binding Ca^2+^, the channels conduct both monovalent cations and phospholipids. Cations permeate without phospholipids, but monovalent cations must occupy the pore for anionic phospholipids to enter. Monovalent cations and anionic phospholipids then permeate the pore in a coupled, electroneutral fashion. Finally, the close coupling of TMEM16F activity and PM expansion leads us to suggest that TMEM16F inactivates as anionic lipids are lost from the cytoplasmic leaflet. In other words, TMEM16F channels, like many ion channels, are activated by binding anionic phospholipids at cytoplasmic regulatory sites (Hilgemann et al., 2018), independent of phospholipid binding within the pore. As to the identity of the membrane compartments that open, caveolae (Shaul and Anderson, 1998) may in principle be involved, consistent with the notion that caveolae open and close in a regulated fashion (Anderson et al., 1992). However, the larger, deeply invaginating components to be described are inconsistent with the morphology of caveolae. From previous work on cell wounding and Ca^2+^ signaling, membrane compartments dubbed “enlargeosomes” in a variety of cell types (Cocucci et al., 2007; Cocucci et al., 2008; Racchetti et al., 2012) are most consistent with our morphological analysis.

## Results

We first describe PM expansion morphometrically and demonstrate the presence of specific VAMP proteins in the compartments that open. Then, we present results relevant to the physiology of PM expansion, including Ca^2+^ requirements, dependence on cell confluence in fibroblasts, and PS exposure in the outer monolayer within deeply invaginating membrane compartments. Next, we describe reversal of PM expansion by extracellular substitution of ions for the disaccharide, trehalose, the activation of PM expansion by cytoplasmic chelation of anionic phospholipid head groups, and the enhancement of PM expansion by sequestering anionic phospholipids in the outer monolayer. Finally, we turn to the molecular function of TMEM16F and demonstrate reversal of PM expansion by manipulating ion gradients and imposing membrane potentials that influence ion distribution inTMEM16F and thereby TMEM16F scrambling function. We provide the composition of 5 standard extracellular (Out) and cytoplasmic (In) solutions in Materials and Methods, and relate any further solution modifications to those 10 solutions.

### Optical characterization of TMEM16F-mediated PM expansion

In a previous study, we used electron microscopy to attempt to identify the membrane compartments in BHK cells that fuse to the cell surface during Ca^2+^ elevations (Yaradanakul et al., 2008). The number of secretory vesicles found within 500 nM of the PM was >10-fold less than would be required to explain the PM expansion observed. Furthermore, PM expansion did not bring ion transporters such as Na/K pumps and constitutively expressed transporters into the PM. As described in the accompanying article (Bricogne, XXXX), analysis by confocal microscopy reveals no involvement of endoplasmic reticulum or acidified lysosomes in PM expansion. As noted already, VAMP2 cleavage by tetanus toxins is without effect on PM expansion (Wang and Hilgemann, 2008; Bricogne, XXXX), and neither super-resolution microscopy nor TIRF microscopy reveals a major involvement of recycling endosomes in PM expansion.

Given these results, it appeared significant that FM dyes reversibly labelled deep membrane compartments during Ca^2+^ elevations (see Movie 1). Since however FM4-64 fluorescence is not proportional to C_m_ (Bricogne, XXXX), we required another dye for quantitative analysis. From several dyes tested (Hilgemann and Fine, 2011) we found that the impermeable dye Trypan blue (TB) fulfilled most of our criteria. TB fluorescence increases when it binds PM sugars and proteins (Harrisson et al., 1981) without insertion into the bilayer as occurs with styryl dyes. As shown in Fig. 2A, TB fluorescence remains proportional to C_m_ during PM expansion and during MEND responses in Jurkat WT and TMEM16F-null cells (Solutions In_1_ and Out_1_, containing physiological monovalent cation concentrations; Pearson correlation coefficient, 0.9; slope, 0.96 ±0.03).

**Figure 2.**
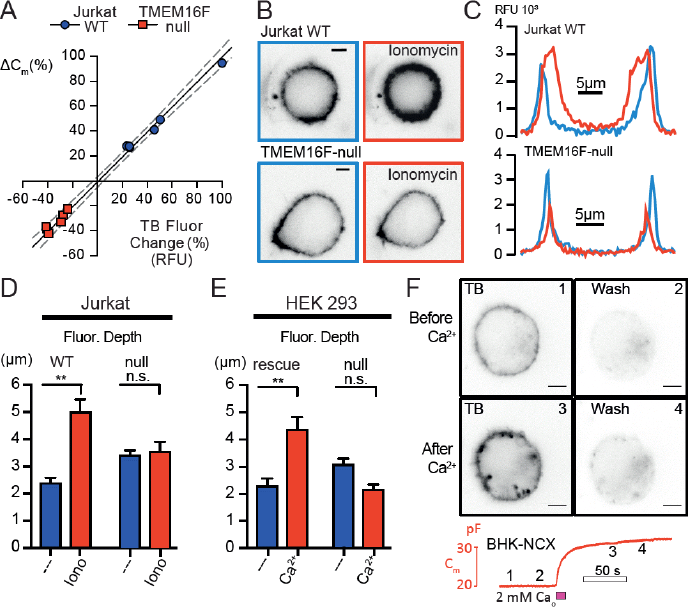
Ionomycin-induced PM expansion involves a membrane compartment that extends deeply into Jurkat, HEK293, and BHK cells. **(A)** Reversible fluorescent membrane staining using trypan blue (TB, 100 µg/ml) was determined before and after application of ionomycin in WT and TMEM16F-null Jurkat cells. Changes of TB fluorescence (%) are plotted against changes of C_m_ (%) caused by ionomycin treatment. Linear regression of results with 95% C.I. confirms a linear correlation between TB fluorescence and membrane area (C_m_) (PCC=0.9). Images **(B)** and fluorescent line scans **(C)** during application of 100 µg/ml TB in Jurkat WT and TMEM16F-null cells before (blue) and after (red) ionomycin-induced PM expansion. Scale bars are 5 µm here and in all subsequent figures. Average membrane fluorescence depth and SEM quantified from **B** for cell lines with and without TMEM16F expression. (**p ≤ 0.01) in Jurkat **(D)** and HEK293 **(E)** cells. Mean values with SEM are given with n>4. **(F)** Confocal imaging of a representative BHK cell, with and without TB application, before and after activation of NCX by 4 mM extracellular Ca^2+^ with 40 mM cytoplasmic Na. The membrane compartments that expand the cell surface extend several microns into the cell and converge to a “neck”-like structure at the interface to the peripheral membrane surface.

Jurkat and HEK293 cells were next patch clamped and labelled with TB, and an initial depth of reversible membrane staining was determined (Figs. 2B-2E, indicated in blue). Cells were subsequently washed, cytoplasmic Ca^2+^ was elevated via ionomycin (5 µM) application, and cells were relabelled with TB for a second analysis (Figs. 2B to 2E, indicated in red). As quantified in Figs. 2D and 2E, the increase in labelling that occurs in WT Jurkat and HEK293 cells after Ca^2+^ elevation extends on average 2.6 and 2.1 µm further into the cells than fluorescence in control measurements. TMEM16F-null cells exhibited no change in labelling depth in response to Ca^2+^ elevations (Fig. 2C). Fig. 2F shows results for NCX1.1-expressing BHK cells in which Ca^2+^ elevations can be induced by activation of reverse Na/Ca exchange (NCX) (Yaradanakul et al., 2008). Similar to HEK293 and Jurkat cells, reversible TB fluorescence also extends many microns deep into the cytoplasm of BHK cells after Ca^2+^ elevations. As evident in the micrographs, individual compartments had remarkably large diameters of >1 µm, appear quite distinct from the PM, and appear to be constricted at their interfaces to the outer PM. Nevertheless, TB labelling rapidly reverses upon washout, thereby verifying connectivity to the extracellular space. Together, these results document that most membrane compartments which open during TMEM16F activation extend deeply into cells and are not morphologically consistent with conventional secretory vesicles, caveolae, or lysosomes. Furthermore, the optical results are consistent with large C_m_ steps (0.1 to 0.5 pF, equivalent to many 100s of secretory vesicles) observed often during PM expansion in fibroblasts (Wang and Hilgemann, 2008).

### Confirmation of VAMP proteins in membrane compartments that open

The morphology of the compartments just described is consistent with reports that novel organelles, dubbed enlargeosomes, expand the cell surface during high Ca^2+^ responses in several cell types (Cocucci et al., 2007; Cocucci et al., 2008). The SNARE protein known as vesicular organelle-associated membrane protein-4, VAMP4, was reported to be enriched in those compartments (Cocucci et al., 2008). Therefore, we expressed VAMP2, 4 and 7 constructs with a luminal pHluorin (pHl) fusion in BHK cells to test whether these proteins undergo a change of accessibility during membrane expansion. The TB absorption spectrum is well suited to quench pHl. Therefore, it is expected that upon compartment opening the exposed fluorescent domains will be quenched by extracellular TB. As shown in Fig. 3 (Solutions In_2_,and Out_2_), TB could indeed be used to determine the initial colocalization of fluorescent signals as well as quenching of new VAMP-pHl fluorescence that appears during Ca^2+^ elevations. To do so, TB was applied and removed both before and after Ca^2+^ elevations. The PM expansion induced by Ca^2+^ influx (see left column of Fig. 3) amounted in each case to ~50%. Before Ca^2+^ elevation, TB (100 µg/ml) caused strong quenching of both VAMP2 and VAMP7 fluorescence, but no quenching of VAMP4 fluorescence (Fig. 3, far right column). Colocalization of TB fluorescence and pHlourin fluorescence (overlaid after TB washout) revealed that VAMP2 has a prominent PM expression which colocalizes with TB fluorescence and quenches 57% of the pHlourin signal (far right column). VAMP7 also has strong surface expression that colocalizes with TB fluorescence. Approximately 50% of VAMP7 pHl signal is quenched by TB. VAMP4 has a distinct intracellular fluorescence pattern with no colocalization with TB fluorescence or quench before Ca^2+^ influx.

**Figure 3.**
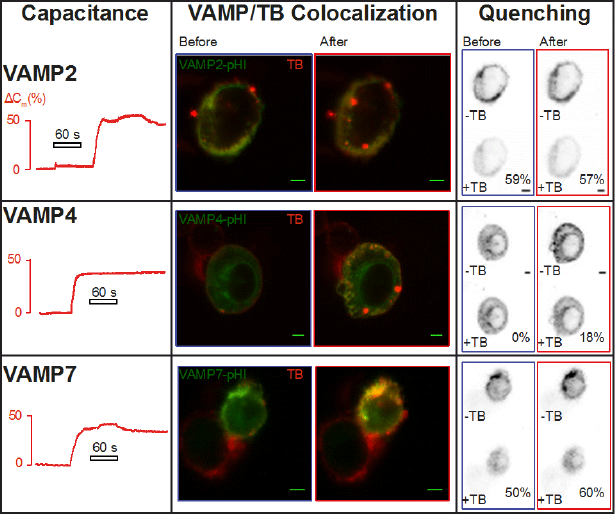
Analysis of VAMP proteins during PM expansion: VAMP4 availability for quenching preferentially increases. Fluorescence recordings in BHK-NCX cells transfected to express green VAMP2, 4, or 7-pHl fusion proteins. Activation of Ca^2+^ influx via NCX causes induces expansion of the PM (left column) corresponding with increases in TB fluorescence (middle column, red). Colocalization is determined by overlay of TB fluorescence and VAMP-pHl after wash off of TB dye. Quenching is demonstrated by the % loss of pHl signal during TB application (right column). VAMP4-pHl fusion protein accessible to the extracellular space, determined by quench of with TB, is negligible in the control state and increases 18% after PM expansion with corresponding increases in colocalization. VAMP2 is already predominantly expressed at the surface and does not increase further after PM expansion while VAMP7 (also present to some extent at the surface) shows moderate increases in colocalization and an increase in TB quenching of 10%. We stress that the VAMP-containing compartments that open, similar to compartments labelled by dyes, appear to be constricted at their interfaces to the outer PM.

Subsequent to PM expansion VAMP2 shows very little change in colocalization and pHluorin quenching, while VAMP7 shows a modest increase in both quenching and colocalization. VAMP7-positive punctae are clearly increased by the Ca^2+^ elevation and are quenched by TB after Ca^2+^ stimulation. VAMP4, however, displays the most robust increase in colocalization and quenching, and the compartments expressing TB-accessible VAMP4 clearly extend deeply into cells. (see Movie 2). We conclude that expressed VAMP4 fusion proteins, and to some extent VAMP7 fusion proteins, accumulate in the membrane compartments that expand the PM. We stress however that these results do not indicate a physical involvement of VAMP proteins in PM expansion but simply support the idea that membrane compartments dubbed enlargeosomes participate in PM expansion, as opposed to endosomes, ER, and lysosomes,

### PM expansion and lipid scrambling with highly controlled free cytoplasmic Ca^2+^

Figures 4 and 5 illustrate experiments using highly Ca^2+^-buffered pipette solutions, rather than Ca^2+^ influx, to activate TMEM16F via a constant free cytoplasmic Ca^2+^ concentration. Fig. 4 shows the time courses of PM expansion and anionic phospholipid exposure in a BHK cell using 4 µM free Ca^2+^. (Solutions In_3_ with 5 mM EGTA and 4.6 mM Ca^2+^ and Out_2_). Membrane area increases by 50% over 10 min, and during this same time the cationic peptide heptalysine-rhodamine (K7r) binds increasingly strongly, but reversibly, to the outer monolayer (Yaradanakul et al., 2008; Bricogne, XXXX). Line scans of the K7r fluorescence illustrate that K7r binding, similar to TB binding, extends deeply into the cell. Movie 3 documents in more detail both the reversibility and depth of K7r binding.

**Figure 4.**
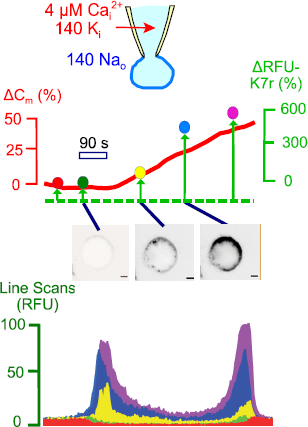
PM expansion and exposure of anionic phospholipids, as monitored by binding of heptalysine-rhodamine (K7r), have similar time courses. A free cytoplasmic Ca^2+^ of 4 µM was maintained by employing highly Ca^2+^-buffered cytoplasmic solutions. Similar to other membrane probes, K7r reveals the opening of deeply invaginating membrane compartments ring PM expansion. Here and in all subsequent figures, concentrations are given in millimolar unless indicated otherwise.

**Figure 5.**
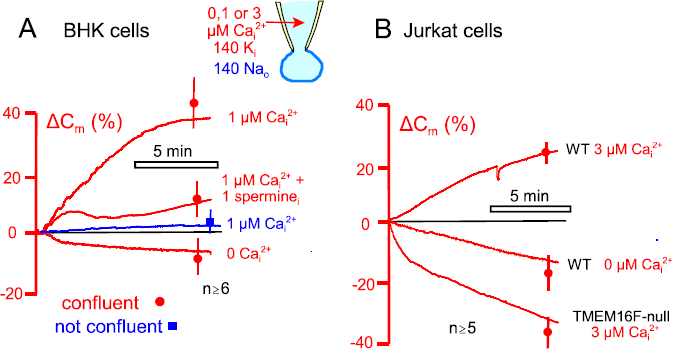
TMEM16F-mediated PM expansion can occur profusely with 1 to 3 µM free cytoplasmic Ca^2+^, albeit slowly. **(A)** In BHK cells that were grown to just confluence (red records and error bars), membrane area (C_m_) increases by 40% over 10 min when cells are perfused with solutions containing 1 µM free Ca^2+^, buffered with 30 mM EGTA. PM expansion does not occur in the absence of Ca^2+^, and PM expansion is blocked by spermine (1 mM), as expected from studies in Jurkat cells. BHK cells removed from dishes during their growth phase (i.e. subconfluent cells, blue record and error bars) exhibit no PM expansion during experiments with 1 µM free cytoplasmic Ca^2+^. **(B)** In Jurkat cells, 3 µM free Ca^2+^ is required to elicit robust PM expansion that amounts to 25% of membrane area over 10 min. Membrane area decreases by 10% over 10 min in the absence of Ca^2+^, and in TMEM16F-null cells membrane area decreases by 35% in the presence of 3 µM free Ca^2+^.

Previously, we described that during acute Ca^2+^ elevations via reverse NCX in BHK cells about 20 µM free Ca^2+^ was required to initiate membrane expansion (Yaradanakul et al., 2008). Given however that NCX currents inactivate and that the Ca^2+^-dependence of membrane expansion shows little or no cooperativity, we re-examined this issue by systematically reducing the free Ca^2+^ to determine the minimal concentrations needed to drive membrane expansion. In presenting these results, we note first for BHK cells that PM expansion can clearly have a strong dependence on the regulatory state of cells. Using EGTA-buffered solutions with cytoplasmic free Ca^2+^ concentrations of 1 to 5 µM, it is our routine observation that PM expansion does not occur when cells are less than 90% confluent (blue experimental record and error bar in Fig. 5A). Using BHK cells that were 90 to 100% confluent, however (red recordings and error bars in Fig. 5A), PM expansion reaches 40% over 12 min when the cytoplasmic solution is buffered to 1 µM free Ca^2+^ (30 mM EGTA, 20 mM Ca^2+^, pH 6.9 with no Mg^2+^ and 2 mM of the nonhydrolyzable ATP analogue, 5’-adenylyl-imidodiphosphate (AMPPNP); Solutions In_4_ and Out_4_). The inclusion of AMPPNP and the absence of Mg^2+^ in cytoplasmic solutions is expected to hinder most ATPase activities. Thus, the results verify that membrane expansion does not require ATPase activities. Employing spermine (1 mM) in the cytoplasmic solutions to block TMEM16F activity (Bricogne, XXXX), PM expansion with 1 µM free Ca^2+^ was decreased by 80%. While this result likely reflects block of the TMEM16F pore by polyamine, it will be described in Fig. 7 that higher concentrations of polyamines can promote significant membrane expansion responses in the absence of Ca^2+^, probably as a result of chelating anionic phospholipids. In further experiments in this same series, PM expansion was completely blocked by removal of cytoplasmic Ca^2+^, and PM area decreased by about 10% over 10 min.

Using Jurkat cells in Fig. 5B, a somewhat higher free Ca^2+^ (3 µM) was required (30 mM EGTA with 27 mM Ca^2+^, pH 7.0) to elicit reliable PM expansion responses of ~25%. For unknown reasons, Jurkat cells lost about 18% of their membrane area over 10 min when perfused with Ca^2+^-free solutions. In TMEM16F-null Jurkat cells, MEND was activated by 3 µM free Ca^2+^ with 30% of the PM being internalized. This is entirely consistent with equivalent experiments using ionomycin in TMEM16F-null Jurkat cells (Bricogne, XXXX). From these results, we conclude that less Ca^2+^ is required for reliable PM expansion than previously estimated, whereby the expansion occurs very slowly with 1 to 3 µM free Ca^2+^. This is indicative of a low affinity Ca^2+^ sensor with a nearly linear dependence on Ca^2+^. Furthermore, the results suggest that PM expansion by the mechanisms analysed here may be subject to physiological regulation.

### PM expansion by TMEM16F activation is reversible

Next, we address a key mechanistic question of this study, namely whether PM expansion indeed reflects fusion of submembrane vesicles with the cell surface. As described in Figs. 6, 8, 12 and 14, PM expansion in BHK cells is readily and rapidly reversible. As background to Fig. 6, disaccharides are known to decrease the lateral mobility of phospholipids in membrane bilayers (van den Bogaart et al., 2007), and trehalose, in particular, is known to affect physical properties and the functionality of lipid bilayers (Kapla et al., 2013). We reasoned therefore that substitution of ions for trehalose might promote membrane-membrane interactions that close the submembrane compartments. When trehalose was substituted for ions in our standard solutions (Solution In_3_ and Out_2_ versus Out_5_), PM expansion induced by reverse NCX activity was decreased within 4 s by 75% of the maximal PM expansion that occurred during Ca^2+^ influx (n=9). Usually, the compartments remained closed after removal of trehalose. The fact that PM expansion can be rapidly reversed suggests that membrane fusion, if it is occurring, is not complete. In this connection, we remind that hypertonic solutions reliably cause exocytosis in most synapses in the absence of Ca^2+^, and this mechanism has been exploited extensively to study exocytosis (Stevens and Williams, 2000). Using similar protocols with hypertonic solutions in BHK and Jurkat T-cells, however, we have never activated significant PM expansion (n>10), suggesting that the PM expansion described here operates by different principles from regulated exocytosis.

**Figure 6.**
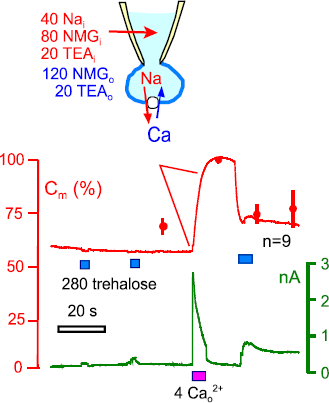
Reversal of PM expansion by replacement of extracellular ions with trehalose. Substitution of ions for trehalose is without any effect on C_m_ before membrane expansion by Ca^2+^ influx (n>10). Ca^2+^ influx via reverse NCX activity results in a 75% increase of C_m_ on average in BHK cells (n=6). Application of trehalose within 30 to 45 s of the expansion response reverses the PM expansion by 86% on average. Thus, membrane compartments that expand the PM during Ca^2+^ influx in BHK cells display a pronounced functional hysteresis, such that they can be reclosed by substitution of extracellular monovalent ions for isoosmotic trehalose.

**Figure 7.**
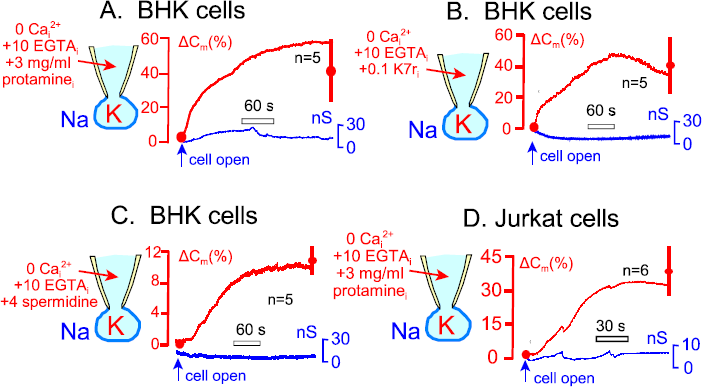
Peptides (heptalysine-rhodamine, K7r) and proteins (protamine) that bind anionic phospholipids cause PM expansion from the cytoplasmic side in the absence of Ca^2+^ (10 mM EGTA). **(A)** BHK cells increase membrane area by 29.7% on average when opened in the cytoplasmic presence of 3 mg/ml protamine. **(B)** BHK cells increase membrane area on average by 37.1 *%* when opened in the presence of 0.2 mM K7r. **(C)** BHK cells increase membrane area on average by 9.4% when opened in the presence of 4 mM spermidine. **(D)** Jurkat cells increase their membrane area by 40.1% on average when opened in the presence of 3 mg/ml protamine.

**Figure 8.**
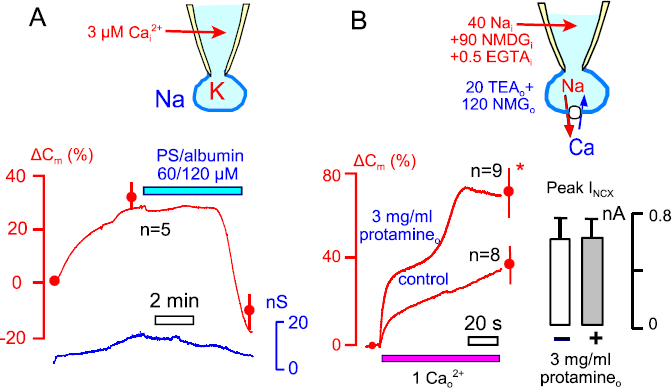
Reversal of PM expansion by extracellular PS-albumin complexes and enhancement of PM expansion by extracellular PS-binding peptides/proteins. **(A)** PM expansion in the presence of 3 µM free cytoplasmic Ca^2+^ can be reversed by extracellular application of PS (60 µM) / albumin (120 µM) complexes. **(B)** PM expansion by reverse NCX activity in BHK cells can be doubled by including cationic proteins that bind PS (protamine, 3 mg/ml) in extracellular solutions. NCX activity is not affected by extracellular application of protamine.

### Polycations activate PM expansion from the cytoplasmic side

We next considered the role of phospholipids in TMEM16F-induced membrane expansion. We hypothesised that anionic phospholipids in the cytoplasmic leaflet could be attachment sites for proteins that hold membrane compartments closed. Accordingly, the scrambling activity of TMEM16F would release the constrictions that close these compartments and open them to the extracellular space. Further, sequestration of anionic phospholipids by cationic peptides, proteins and other polycations should in principle release the constrictions and trigger membrane expansion. In the experiments described in Fig. 7, the cytoplasmic solution contained 10 mM EGTA with no added Ca^2+^, no added Mg^2+^, no ATP, and no GTP (Solutions In_5_ and Out_4_). As shown in Fig. 7A, C_m_ increased over 5 min by 40% when BHK cells were opened in the presence of 3 mg/ml protamine, a cationic protein that binds avidly negatively charged phospholipids of biological membranes (Ichev et al., 1985) and reverses the anticoagulant effects of heparin (Carr and Silverman, 1999). As shown in Fig. 7B, C_m_ increased over 5 min by 37% when BHK cells were opened in the presence of 0.1 mM K7r in the cytoplasmic solution. As shown in Fig. 7C, a high concentration of spermidine (4 mM), higher than required to block membrane expansion and promote fast MEND responses (Lariccia et al., 2011), was partially effective in BHK cells, increasing C_m_ by ~10%. Finally, as shown in Fig. 7D, C_m_ increased by 37% over 2 min when Jurkat cells were opened in the presence of 3 mg/ml protamine.

### PS gradients can drive forward and reverse PM expansion

The experiments just described lend initial support to the idea that loss of negatively charged lipids from the cytoplasmic membrane leaflet, rather than a gain of negatively charged lipids in the outer monolayer, mediates the opening of membrane compartments. Further key predictions are that, when TMEM16F is active, an enrichment of PS in the extracellular monolayer should promote compartment closure, while sequestration of PS on the extracellular side should shift PS from the cytoplasmic side to the extracellular side and thereby promote PM expansion.

As outlined in Fig. 8A, we enriched cells with PS by applying PS-albumin complexes prepared in a 1:2 ratio (Bartholow and Geyer, 1982) by sonication of natural PS from bovine brain with albumin in distilled water. Using cytoplasmic solutions with 3 µM free cytoplasmic Ca^2+^ (20 mM EGTA with18 mM Ca^2+^), PM expansion amounted to 35% on average in 5 experiments. Application of 120 µM albumin with 60 µM PS then resulted in complete reversal of PM expansion after a delay of 3 to 4 min. In fact, C_m_ values decreased significantly below baseline, as expected if PS activates significant endocytic responses in addition to reversing PM expansion. We note that other anionic phospholipids, especially PIP_2_, can promote endocytic processes when perfused into cells (Lariccia et al., 2011).

Figure 8B demonstrates that sequestration of anionic phospholipids on the extracellular side promotes PM expansion. In these experiments, we used Na/Ca exchangers to generate Ca^2+^ influx in BHK cells. We then compared PM expansion in the presence and the absence of 3 mg/mL extracellular protamine (Solution In_3_ and Out_2_). PM expansion during application of 1 mM extracellular Ca^2+^ for 2 min amounted to 40% and was increased to 80% (p=0.02) in the presence of extracellular protamine. Importantly, peak NCX currents were unchanged. In summary, extracellular PS can reverse Ca^2+^ -induced PM expansion in BHK cells, albeit with a delay, while sequestration of extracellular PS can promote PM expansion as expected if PS is in rapid equilibrium between monolayers during Ca^2+^ elevations.

### Relationships between TMEM16F conductance and PM expansion

We analyze next the specific relationships between monovalent cation concentrations, ion permeation, and PM expansion during TMEM16F activation. Fig. 9 describes results for Jurkat cells in which PM expansion was monitored via capacitance (C_m_) changes, and membrane current (I_m_) and conductance (G_m_) were monitored in parallel. Ca^2+^ elevations were induced by ionomycin (5 µM) application in the presence of 2 mM Ca^2+^ for up to 1 min. As shown in Fig. 9A (Solutions In_1_ and Out_1_), the time course with which C_m_ rises follows closely the time course of membrane current (I_m_) and conductance (G_m_). Specifically, the integral of the G_m_ signal corresponds to the time course of the C_m_ signal, after scaling, and the derivative of the C_m_ signal corresponds to the time course of the G_m_ signal after scaling (>20 observations). Given this close relationship, we next tested whether TMEM16F conduction of ions per se controls PM expansion. To do so, we replaced cytoplasmic and extracellular ions with aspartate (Asp=As), *N*-methyl-d-glucamine (NMG=Nm), or isoosmotic sucrose (Suc) (Fig. 9B; Solutions In_3_ and Out_2_). The presence of either cytoplasmic Na (40 mM in Fig. 9B) or K supported large-scale PM expansion in the absence of Cl, while expansion responses were smaller without Na or K on the cytoplasmic side but with NaCl outside. Responses were reduced by ~50% from control responses when Na and K were replaced with equimolar NMG-Asp on both membrane sides, and they were reduced by 80% when TEA-As was employed on both membrane sides. Replacement of 90% of ions with isotonic sucrose fully blocked exocytosis.

**Figure 9.**
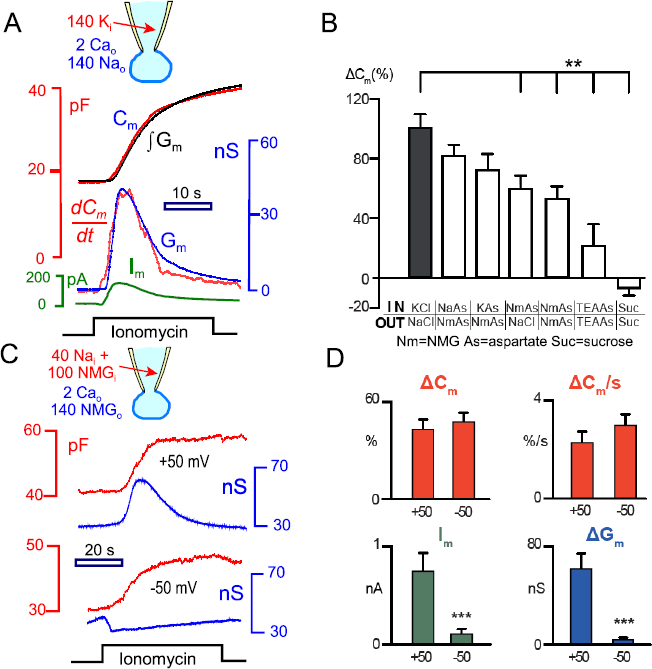
Relationships between membrane conductance (G_m_), current (I_m_) and capacitance (C_m_) during ionomycin-induced PM expansion in Jurkat T-cells. **(A)** Using standard experimental solutions with WT Jurkat cells, G_m_ and I_m_ have similar time courses (lower recordings), and the integral of G_m_, ∫G_m_, tracks the time course of PM expansion (ΔC_m_) (upper recordings) during application of ionomycin (5 µM). Records were scaled to start and end at the same points, n >10. **(B)** Percentage change in C_m_ 30 s after ionomycin treatment of WT Jurkat cells. Cells were perfused with indicated cytoplasmic solutions for >200 s before ionomycin treatment. Extracellular (Out) and cytoplasmic (In) solutions were as shown: Standard cytoplasmic (KCl), Sodium Aspartate (NaAs), Potassium Aspartate (KAs), NMG Aspartate (NmAs), Ringer solution (NaCl), TEA and isotonic sucrose (Suc) with 0.5 mM EGTA with 4 mM (Out) or 0.5 mM MgCl_2_ (In). **(C and D)** Ion permeation does not influence PM expansion. Solutions were symmetrical NmAs with 30 mM NMG replaced by 30 mM Na on the cytoplasmic side to generate outward Na current. Membrane potential was held at +50 or –50 mV during the entire experiments. Representative recordings are shown in C; mean values and SEM for n=7 at each potential are shown in **D.** Ionomycin treatment transiently increases conductance (blue) and generates outward Na current (green) at +50 mV but not at -50 mV. However, C_m_ (red) increases to the same extent and at the same maximal rate at -50 and +50 mV.

The relative changes of PM expansion with different ions correspond roughly to the relative TMEM16F permeabilities to Na, K, NMG, and TEA (Yang et al., 2012). Therefore, we tested whether PM expansion is influenced by ion permeation per se. Fig. 9C illustrates experiments in which we employed an outwardly directed Na gradient (30 mM cytoplasmic Na with 110 mM cytoplasmic NMG and 140 mM extracellular NMG), so that TMEM16F current became strongly voltage dependent. From the experiments of Fig. 9B, it may be assumed that PM expansion in this condition is limited in rate and extent by the cytoplasmic Na concentration. In contrast to results with equal K and Na concentration on opposite membrane sides (Fig.9A), large (>0.5 nA) outward Na currents are generated in these experiments (Fig. 9D green). Holding membrane potential at +50 mV, ionomycin causes a large transient conductance, as in Fig. 9A, which develops and decays with the same time course as PM expansion (Fig.9C top and Fig. 9D blue). When membrane potential is held at -50 mV, both the conductance and outward membrane currents are strongly suppressed. Nevertheless, PM expansion proceeds to the same extent (ΔC_m_) and with the same maximal rate (ΔC_m_/dt) as at +50 mV (Fig. 9C-D red). Composite results (n=6) shown in Fig. 9D therefore dissociate ion conduction *per se* from PM expansion. The results also negate a possibility that Ca^2+^ influx via TMEM16F initiates PM expansion because, in that case, hyperpolarization would enhance Ca^2+^ influx and thereby PM expansion. Rather, another function of TMEM16F must underlie the activation of PM expansion.

### Anionic phospholipid scrambling and PM expansion require monovalent cations

As outlined in the Introduction, our further experiments suggest that monovalent cations must occupy the TMEM16F pore before anionic phospholipids can enter and permeate the pore (see Fig. 1). The actual permeation of anionic phospholipids then takes place in an electroneutral fashion that is not influenced by membrane voltage. To explain this suggestion, we provide in Fig. 10 simple simulations of the working hypotheses, assuming for simplicity that TEA cannot occupy the TMEM16F pore and that Na, K and Cs occupy the pore with equal affinity. Simulation details are provided in Material and Methods. Each black curve in the Fig. 10 depicts the fractions of invaginating membrane compartments predicted to be open (F_open_) as a result of anionic phospholipids being lost from the cytoplasmic to the extracellular leaflet. It is assumed that 4 phosphatidylserine (PS) molecules are bound by membrane proteins to close the compartments, and it is assumed that PS binding is half-saturated at approximately the resting concentration of PS in the inner monolayer (i.e. with100 arbitrary PS units). Here and for subsequent simulation results, we assume that PS has a 4-fold higher affinity for the outer monolayer than for the inner monolayer. This assumption reflects the fact that the outer monolayer is more ordered than the inner monolayer, resulting in 5-times slower diffusion of phospholipids in the outer monolayer than in the inner monolayers (Cribier et al., 1990). As a result, the majority of PS translocates to the outer monolayer during TMEM16F activation.

**Figure 10.**
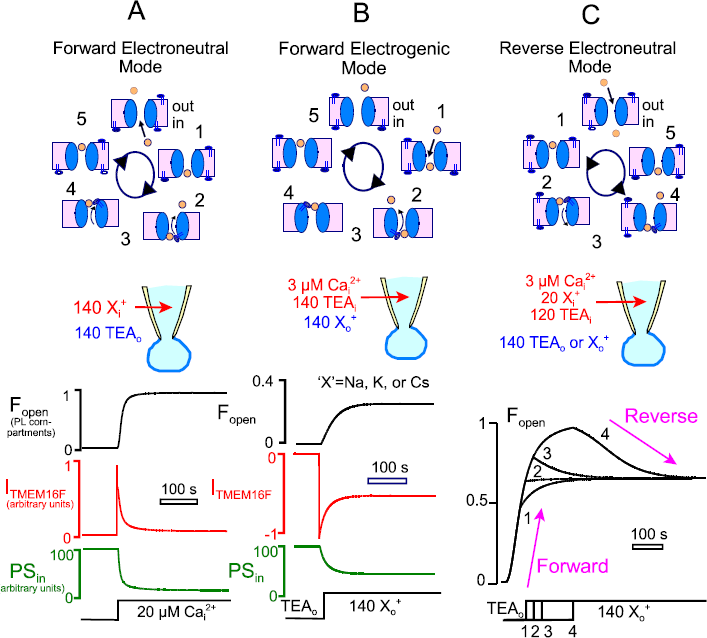
Simulation of TMEM16F function and PM expansion accompanying TMEM16F activation. **(A)** Predicted results for activation of TMEM16F by increasing cytoplasmic free Ca^2+^ from 0 to 20 µM (similar to reverse NCX activation or ionomycin exposure). Nearly all subsurface PM compartments open to the extracellular space, TMEM16F current inactivates by about 90%, and >80% of PS translocates from the cytoplasmic leaflet to the extracellular membrane leaflet. **(B)** PM expansion induced by applying 140 mM permeant cation on the extracellular side in the absence of permeant cations on the cytoplasmic side. PS translocates to the outside in an electrogenic fashion when a cation permeates inwardly and then retreats back to the extracellular side together with one PS molecule. **(C)** PM expansion with 140 mM extracellular TEA and a cytoplasmic solutions containing TEA and 20 mM permeant cation. Responses 1 to 4 were generated by substituting 140 mM extracellular TEA for permeant cations at progressively later time points. When PM expansion approaches steady state (4), PM expansion reverses by 30% when extracellular TEA is replaced by permeant cations.

Three simulations are presented in Fig. 10, one with permeant cations only on the inside (A), one with permeant cations only on the outside (B), and one upon changing monovalent cation gradients from outward to inward (C). In Fig. 10A the cytoplasm is assumed to contain 140 mM of monovalent cation (X) while the extracellular solution contains 140 TEA (i.e. no TMEM16F-interacting cations). Upon increasing cytoplasmic free Ca^2+^ in a step from 0 to 20 µM, the fractional opening of invaginating membrane compartments (F_open_) increases from <0.1 to nearly 1 within 10 s, TMEM16F generates an outward current that inactivates as PS is lost from the cytoplasmic side, and the amount of PS in the cytoplasmic leaflet decreases from nearly 100% of the PM PS to about 15%. The cartoon above the simulation illustrates the partial reactions that scramble PS under these conditions and that occur in an electroneutral fashion. When a cation binds to the cytoplasmic end of the pore (#1), a PS molecule can bind from the cytoplasmic side (#2), the cation and PS can traverse the pore (#3), PS can move from the pore to the outer monolayer (#4), and the cation can exit the pore (#5). As illustrated in Fig. 10B, TMEM16F according to our model cannot scramble anionic phospholipids in the absence of permeable monovalent cations. When permeable extracellular cations are applied, an inward cation current is generated. As cations reach the inner mouth of the pore, they support the movement of PS to the extracellular side and return to the extracellular side with PS. As illustrated in the cartoon above the simulation, the cycle of reactions moves one negative charge outwardly per cycle, and this reaction cycle can therefore be activated by hyperpolarization. Fig. 10C illustrates the induction of reverse PS scrambling. Initially, it is assumed here that the cytoplasm contains 20 mM permeant cations while the extracellular solution contains no permeant cations. Over time, the majority of PS reaches the extracellular monolayer and accumulates in the outer monolayer as a result of its higher affinity for PS than the inner monolayer. When a concentration of 140 mM permeant cations is applied to the outside at an early time point, scrambling is stabilized at an intermediate level. If PS is allowed to accumulate further in the outer monolayer, application of extracellular permeant cations can partially reverse scrambling via the reaction sequence illustrated in the cartoon for reverse electroneutral transport.

Figures 11 and 12 verify each of these model predictions by experiments. Fig. 11 shows the activation of scrambling and PM expansion when experiments are initiated without permeant cations on the cytoplasmic side. A free cytoplasmic Ca^2+^ concentration of 5 µM was employed with, at first, 140 mM TEA on both membrane sides. PS exposure was monitored via periodic application of K7r and measurement of its membrane-associated fluorescence. After 3 min in the presence of TEA on both membrane sides, C_m_ remains stable and the K7r signal remains small and stable. After substituting extracellular TEA for Cs (140 mM; Solution Out_3_), however, C_m_ begins to rise with a delay of about 30 s, and total PM expansion amounts to 80% at 10 min. The reversible K7r signal grows roughly in proportion to membrane area, and reveals reversible binding of K7r into compartments that extend multiple microns into the cell (see micrograph #3).

**Figure 11.**
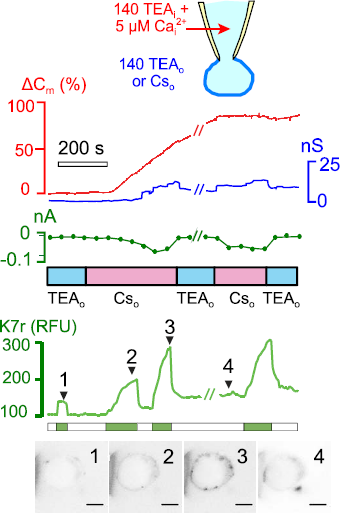
Activation of PM expansion and PS scrambling to the outer monolayer by extracellular Cs. The experiment was initiated with TEA on both membrane sides and with 5 µM Ca^2+^ on the cytoplasmic side. C_m_ is stable and K7r binding is initially very low. After 2.5 min, extracellular TEA was substituted for Cs, and PM expansion began within 1 min. K7r binding increases in parallel with PM expansion and remains reversible. Deep membrane compartments become labeled over time (see micrograph #3).

**Figure 12.**
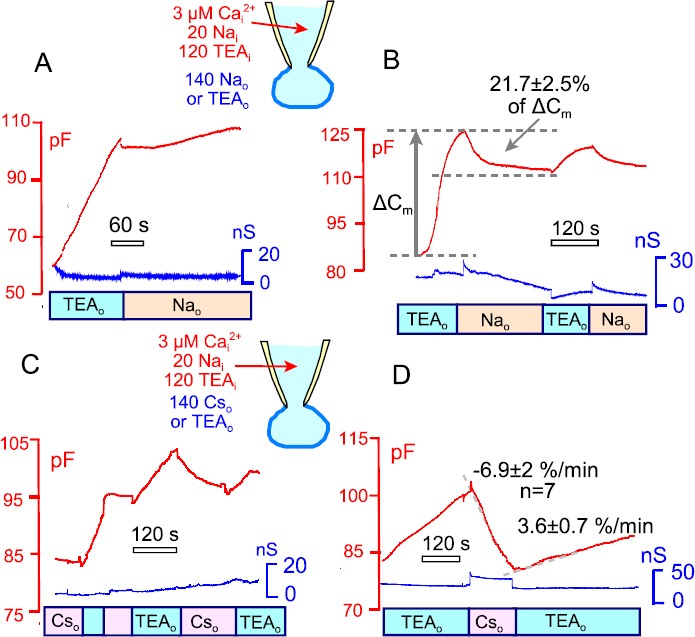
As predicted from simulations, experiments reveal inhibition and reversal of PM expansion by 140 mM extracellular Na and Cs in the presence of 20 mM cytoplasmic Na and 3 µM cytoplasmic free Ca^2+^. **(A)** PM expansion proceeds rapidly in the presence of extracellular TEA, and the expansion stops when cytoplasmic TEA is substituted for Na. **(B)** After PM expansion reaches a near steady state in the presence of extracellular TEA, PM expansion reverses by 22% on average when extracellular TEA is substituted for Na. **(C)** Blockade and partial reversal of PM expansion by substituting extracellular TEA for Cs. **(D)** Reversal of ongoing PM expansion upon substituting extracellular TEA for Cs. The rate of membrane area change adjusts within just a few seconds upon changing ion gradients.

Figure 12 verifies further predictions of simulations by experiments in which either extracellular Na (Figs. 12A and 12B) or extracellular Cs (Figs. 12C and 12D) were used to slow and/or reverse PM expansion (Solutions In_5_ and Out_4_). Cytoplasmic free Ca^2+^ was heavily buffered with 30 mM EGTA to 3 µM (30 mM EGTA/27 mM Ca^2+^). Fig. 12A illustrates the case in which PM expansion is initiated with a low cytoplasmic cation concentration (20 mM Na) with 140 mM extracellular TEA. Substitution of TEA for 140 mM Na on the outside effectively stops PM expansion, as in the 2^nd^ simulation in Fig. 10C. Fig. 12B illustrates records in which PM expansion is allowed to proceed to nearly its maximal extent with 3 µM free Ca^2+^ (as in curve #4 in Fig. 10C). In this case, PM expansion reverses by about 25% upon substituting extracellular TEA for Na, and expansion can be reactivated by the reverse concentration change. Using Cs as a substitute for TEA in Fig. 12C, Cs initially stops PM expansion at an intermediate value, as in Fig. 12A with Na. After expansion is allowed to proceed further in the presence of TEA, Cs causes an approximately 25% reversal of PM expansion, equivalent to 10% of total membrane area, similar to results for extracellular Na in Fig. 12B. Fig. 12D illustrates in more detail the speed with which membrane area changes can be reversed by applying and removing Cs in these experiments. The maximal rates of membrane area change become established within just a few seconds after changing the extracellular solution composition. We mention that, in individual experiments, the rates of membrane area change required up to 30 s to change after changing solutions, perhaps because intracellular cation concentrations can change significantly during these experiments and require time to readjust after changing concentration.

### PM expansion can be reversed by depolarization

Finally, we consider effects of membrane potential on PS scrambling and PM expansion with the aid of simulations. With physiological ion concentrations on both membrane sides, simulations predict no effect of membrane potential because the TMEM16F channel is nearly saturated with cations. This is illustrated in Fig. 13A (140 mM monovalent cations on both membrane sides; dissociation constant, 15 mM at both channel orifices). Voltage changes from -70 to +70 mV have no effect on PS scrambling or compartment opening, as described experimentally in Fig. 9A. Fig. 13B illustrates simulations with 40 mM permeant cation on the cytoplasmic side and no permeant cation on the outside, equivalent to experiments presented in Fig. 9B. Voltage changes from -70 to +70 mV are nearly without effect because cytoplasmic pore sites are nearly saturated. The lack of effect of membrane voltage is additionally supported by the assumption that PS binds with higher affinity in the extracellular than in the cytoplasmic monolayer. Owing to this asymmetry, voltage must be increased to >120 mV to effectively drive an electrogenic reverse flux of PS.

**Figure 13.**
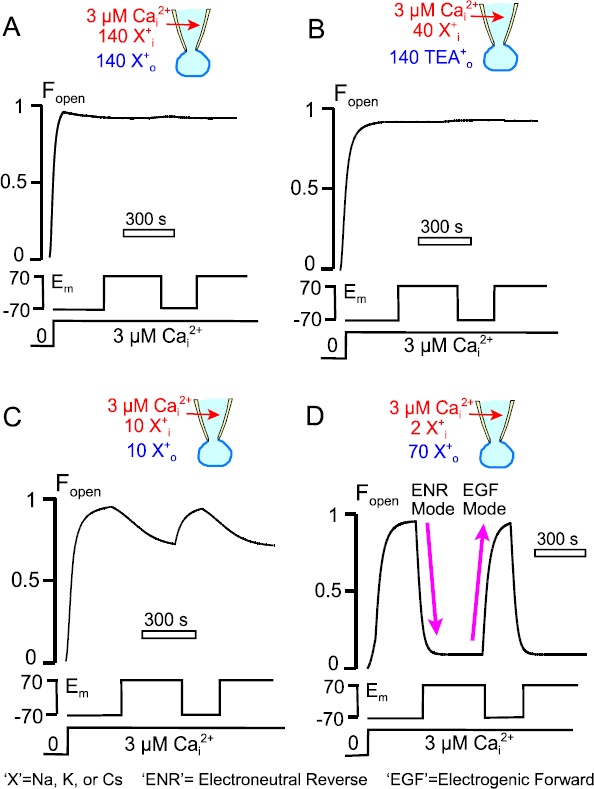
Modeled effects of membrane potential with different permeant cation concentrations and 3 µM cytoplasmic free Ca^2+^. In A, B, C, and D, the fraction of open invaginations (F_open_) is simulated with 140 mM permeant cations on both membrane sides **(A),** as in Fig. 9A, with 40 mM permeant cations on the inside and no permeant cations on the outside **(B)**, as in Fig. 9C, with 10 mM permeant cations on both sides **(C),** and with 2 mM permeant cations on the inside and **70** mM permeant cations on the outside **(D)**, as in Fig. 14 Membrane potential changes are without effect in A and B because cation binding does not limit PS translocation. With low cation concentrations (C), depolarization favors an outward cation gradient and thereby outward PS movement. However, the high affinity of PS for the outer monolayer limits the influence of membrane potential within our experimental range. In D, the large inward gradient of permeant cations overcomes the PS balance to the outer monolayer and promotes reverse PS transport via electroneutral reverse (ENR) transport. Still, to activate substantive reverse PS transport, the membrane must be depolarized to disable electrogenic forward (EGF) transport of PS. When the membrane is hyperpolarized, EGF PS transport rapidly reopens the membrane compartments.

**Figure 14.**
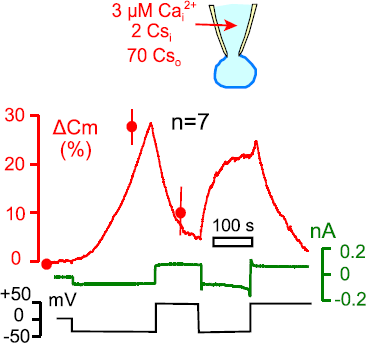
In the presence of a large inward gradient of permeant monovalent cations, PM expansion is promoted by hyperpolarization and reversed by depolarization. The cytoplasmic solution contains no ATP, no Mg^2+^ and no GTP. Using 2 mM cytoplasmic and 70 mM extracellular Cs in the presence of additionally 138 and 70 mM TEA, respectively, hyperpolarization drives extensive PM expansion that amounts to 30% over 4 min. Upon depolarization, PM expansion reverses, and responses can be repeated. Hyperpolarization favors an electrogenic outward movement of PS under these conditions, while depolarization stops the electrogenic PS transport mode and allows reverse PS transport from the outer to the inner monolayer. See Fig. 13D for comparison to model predictions.

As expected, membrane potential effects become significant when permeant cation concentrations are reduced to the assumed dissociation constants for ion binding that initiates PS binding. As shown in Fig. 13C with 10 mM permeant monovalent cations on both membrane sides, the fraction of open compartments (F_open_) increases and decreases about 20% during hyperpolarization to -70 mV versus depolarization to +70 mV. These effects are still not particularly large, and Fig. 13D illustrates the optimal conditions for effects of membrane voltage. In brief, a large inward cation gradient is required. Specifically, the presence of a low concentration of permeant cations (2 mM) in the cytoplasm and a high cation concentration (70 mM) on the extracellular side overcomes the tendency of PS to accumulate preferentially in the outer monolayer when TMEM16f channels are open. Hyperpolarization then strongly favors electrogenic forward PS transport, and therefore compartment opening, while depolarization negates the electrogenic mode and allows efficient electroneutral reverse PS permeation, thereby favouring closure of the PS-dependent membrane compartments.

Figure 14 presents the equivalent experiments employing the conditions just outlined (5 mM cytoplasmic Cs with 70 mM extracellular Cs). We included 2 mM AMPPNP in the cytoplasmic solutions to block ATPase activities. Using 5 mM EGTA with 4.5 mM Ca^2+^ (~4 µM free Ca^2+^), membrane area expands in a progressively accelerating fashion at -30 mV and reaches 130% of control area after 5 min. Upon switching membrane potential to +50 mV, membrane area immediately begins to decline at nearly the opposite rate (~30% per minute) and declines nearly to baseline. In all experiments, these responses could be repeated at least once, and often multiple times, to nearly the same extent as the initial responses. The reversible nature of PM expansion in these experiments indicates that membrane fusion events, if they are occurring, are reversible. Alternatively, the opening and closing of invaginating membrane compartments might not involve membrane fusion and excision events but rather the opening and closing of constrictions that are adequate to isolate them electrically from the extracellular space.

## Discussion

It has long been known that diverse cell types can expand their PM by substantial amounts in response to large elevations of cytoplasmic Ca^2+^ (Coorssen et al., 1996; Ninomiya et al., 1996; Kasai et al., 1999; Togo et al., 2003; Kreft et al., 2004; Bretscher, 2008; Cohen et al., 2008; Yaradanakul et al., 2008). This expansion has often been suggested to mediate repair of membrane wounds (Togo et al., 2003; Bradke et al., 2012; Cooper and McNeil, 2015; Corrotte et al., 2015), and an involvement of SNARE-mediated mechanisms seems well established in some cell wounding protocols (Bi et al., 1995). In a companion article (Bricogne, XXXX), we have shown that TMEM16F can initiate large-scale PM expansion in multiple cell types. Here we have demonstrated that this type of PM expansion is kinetically and mechanistically coupled to scrambling of anionic phospholipids, which in turn requires occupation of the TMEM16F pore by monovalent cations. Clearly, the underlying mechanisms are substantially different from classical Ca^2+^-activated exocytosis. PM expansion described here reflects to a large extent the opening of deeply invaginating PM compartments, and the openings appear to be regulated by constrictions in junctional regions close to the cell surface.

### PM expansion may not involve *bona fide* membrane fusion events

Our results show a very close correlation between the opening of TMEM16F channels and the expansion of the PM in Jurkat cells (Figure 9). The time course of TMEM16F conductance changes matches closely the time course of PM expansion. Specifically, the integral of conductance signals matches the rising capacitance waveform, as if TMEM16F channel activity directly controls PM expansion. Ironically, genuine exocytic membrane fusion events generate a transient pseudo-conductance in electrical recordings that reflects the expansion of so-called fusion pores (Lindau, 2012). We stress therefore that the conductance signals monitored in our experiments via square-wave voltage perturbation reflect a true PM conductance that follows the rise and fall of membrane current during membrane expansion (Figure 9A).

Although cytoplasmic monovalent cations are required for PM expansion (Figures 9 and 11), ion conduction per se can be entirely dissociated from PM expansion by changing the membrane holding potential (Figures 9C and 9D). Rather, it appears that monovalent cations serve as counter charges for the movement of anionic phospholipids through TMEM16F. This requirement would be rationally explained by the existence of negative charges in the TMEM16F pore that facilitate cation conduction while, at least to some extent, repelling anions and especially anionic phospholipids. The parallel block of TMEM16F conductance and PM expansion by spermine ((Bricogne, XXXX); Fig. 3), possibly as a result of polyamine binding within the negatively charged pore, is also consistent with lipids and ions moving through the TMEM16F channel in a concerted fashion. To what extent polyamines also inhibit TMEM16F activity by sequestering anionic phospholipids into nanodomains (Denisov et al., 1998), and thereby promoting TMEM16F inactivation, is not certain. At higher concentrations than are required to block TMEM16F activity, polyamines can promote membrane expansion in the absence of Ca^2+^ (Figure 7), probably by binding anionic phospholipid head groups in a manner similar to K7r and protamine.

The similar time courses with which the PM expands and extracellular K7r binding capacity increases (Figures 4 and 11) are suggestive of a simple relationship between PS translocation and compartment opening. This impression may be to some extent fortuitous because K7r will simultaneously bind multiple anionic phospholipids (Denisov et al., 1998) and, as represented in our simulations, closure of the membrane compartments likely requires binding of multiple anionic phospholipids by membrane-interacting proteins. Most importantly, we have documented that PM expansion caused by activation of TMEM16F can be partially or completely reversed by establishing large inward cation gradients and by membrane depolarization (Figures 9 and 11). These results seem inconsistent with the idea that PM expansion is occurring via genuine membrane fusion events. Either the fusion events are incomplete and impeded at some step, or else there are no fusion events. The major possibility that emerges is that PM expansion during TMEM16F activation involves membrane that is contiguous with the PM yet uncoupled electrically from the extracellular space. As PS is depleted from the inner monolayer, constrictions in junctional regions of these compartments relax and allow access to the extracellular space.

### The membrane compartment that opens

The hypothesis that PM expansion reflects relaxation of membrane binding proteins that constrict the necks of these compartments is supported by the following evidence: (1) PM expansion is reversible as expected for a direct dependence on protein binding to anionic phospholipids (Figures. 9, 11, and 13). (2) Large-scale PM expansion can be initiated in the complete absence of cytoplasmic Ca^2+^ by ‘sequestering’ anionic phospholipids with cytoplasmic peptides or proteins that bind their head groups with substantial affinity (Figure 12). (3) PM expansion can be reversed by increasing the PS content of the PM via application of PS-albumin complexes from the extracellular space, and this occurs specifically under conditions in which TMEM16F activity should be substantial. (4) PM expansion is enhanced by incubating cells with proteins, e.g. protamine, or peptides, K7r, that bind anionic phospholipids and therefore will shift the equilibrium of PS from the cytoplasmic to the extracellular PM monolayer (Figure 13).

The membrane that expands the PM is clearly not a classical membrane compartment. As stressed throughout this article, expansion involves extended, deeply invaginating membranes that can have complex dimensions extending over several microns (Figures 2 and 3). Extended membrane networks, especially in close proximity to the cell surface, might be confused with ER in electron microscopy studies. We cannot exclude at this time that caveolae become involved in PM expansion, and it is well established that the assembly of caveolae is highly dependent on PS (Hirama et al., 2017). Experiments described in the companion article (Bricogne, XXXX) limit strongly potential roles for recycling endosomes and lysosomes, and a role for endoplasmic reticulum is effectively eliminated. The appearance of the vesicle membrane protein, VAMP4, in the PM during PM expansion confirms a previous report (Cocucci et al., 2008). This does not however add substantively to an understanding of this compartment or for that matter the role of SNARE proteins in PM expansion. Importantly, the membrane that opens contains few if any PM ion transporters or ion channels that are otherwise abundant in the PM (Yaradanakul et al., 2008). Finally, we stress that our experiments do not address whether PM expansion, as described here, might play a role in membrane repair. The complete relaxation of constrictions would presumably allow the compartments to rapidly merge with the pre-existing PM, as would be required for wound repair.

### Key problems and questions raised by constriction-dependent PM expansion

This study does not address the identity of the membrane proteins that mediate membrane constriction. Obvious candidates include (1) dynamins (Schmid et al., 1998; Antonny et al., 2016), (2) BAR domains (McMahon and Boucrot, 2015; Suetsugu, 2016; Salzer et al., 2017), and (3) dysferlins (Demonbreun and McNally, 2016). For now, we have no reason to favour one over another. Membrane expansion can occur with as little as 1 µM free Ca^2+^ under control of TMEM16F phospholipid scrambling (Figure 5), and each of these membrane-shaping systems could potentially regulate membrane expansion via additional control mechanisms.

A second key problem is the biogenesis of the invaginations that open during PM expansion. Compartment growth might occur via trafficking and fusion of endosomes to pre-existing structures at the PM, and our TIRF analysis suggests that recycling endosomes can contribute to some extent (Bricogne, XXXX). Compartment growth might also occur via the slow internalization of surface bilayer through the restrictions that hold the compartments closed. The passage of integral membrane proteins into the compartments would presumably be restricted, thereby explaining the relative lack of typical PM transporters and channels in the compartments that open (Yaradanakul et al., 2008).

Finally, it will be of substantial interest to test whether similar membrane compartments exist in cells that do not express TMEM16F, for example by testing whether polycations can open membrane compartments in candidate cells from the cytoplasmic side. When this is known, it will then be more readily possible to determine which physiological cell processes are functionally dependent on this unexpected form of PM expansion. The list may in principle include all physiological functions attributed to conventional exocytosis. That is to say, this form of PM expansion may contribute membrane to close cell wounds, as well as to allow cell growth and swelling. It might also release cytokines and other chemicals to the extracellular space, and it may bring specific membrane proteins into the cell surface in a regulated fashion.

## Acknowledgments

We thank Mei-Jung Lin for technical help and discussion. CB was supported by The Rosetrees Trust (M155), MRC Doctoral Training Grant (1132770), and the UCL Bogue Fellowship. DH and MF were supported by National Institutes of Health, USA (HL119843, T32DK007257).

## Materials and Methods

Methods were as in the accompanying article (Bricogne, XXXX). Baby hamster kidney (BHK) cells expressing cardiac NCX1.1 were grown as described (Yaradanakul et al., 2008). Cells were imaged using the Nikon EZ-C1 confocal microscopy system as described in the companion article (Bricogne, XXXX) and in the text. Fluorescent line scans were determined by subtracting the equivalent background fluorescence. Membrane fluorescence depth in microns was determined from line scan images and quantified in ImageJ using 4 distinct membrane regions per cell.

### Solutions

Table 1 provides the compositions of major solutions employed. Further solution modifications are given with reference to these solutions in Results.

**Table 1.**

| **Solutions**<br>**(mM)** | **Cytoplasmic ("In")** |  |  |  |  | **Extracellular ("Out")** |  |  |  |  |
| --- | --- | --- | --- | --- | --- | --- | --- | --- | --- | --- |
|  | **#1** | **#2** | **#3** | **#4** | **#5** | **#1** | **#2** | **#3** | **#4** | **#5** |
| Dextrose | 0 | 0 | 0 | 0 | 0 | 10 | 0 | 0 | 0 | 0 |
| HEPES | 10 | 20 | 20 | 30 | 20 | 10 | 10 | 10 | 10 | 10 |
| KOH | 140 | 0 | 140 | 160 | 0 | 5 | 0 | 0 | 0 | 0 |
| NaOH | 5 | 40 | 0 | 0 | 20 | 130 | 0 | 0 | 0 | 0 |
| CsOH | 0 | 0 | 0 | 0 | 0 | 0 | 0 | 140 | 0 | 0 |
| TEA-OH | 0 | 20 | 0 | 0 | 120 | 0 | 20 | 0 | 140 | 0 |
| NMG | 0 | 80 | 0 | 0 | 0 | 0 | 120 | 0 | 0 | 0 |
| EGTA | 0.5 | 0.5 | 5 | 30 | 30 | 0 | 0.5 | 0.5 | 0.5 | 0.5 |
| CaCO<sub>3</sub> | 0 | 0 | 0 | 20 | 25 | 0 | 0 | 0 | 0 | 0 |
| CaCl<sub>2</sub> | 0.25 | 0.25 | 4.6 | 0 | 0 | 0 | 0 | 0 | 0 | 0 |
| MgCl<sub>2</sub> | 0.5 | 0.5 | 0.5 | 0 | 0 | 4 | 4 | 4 | 4 | 4 |
| Aspartate | 0 | 115 | 125 | 90 | 125 | 0 | 125 | 125 | 125 | 0 |
| HCl | 145 | 0 | 0 | 0 | 0 | 135 | 0 | 0 | 0 | 0 |
| Trehalose | 0 | 0 | 0 | 0 | 0 | 0 | 0 | 0 | 0 | 300 |
| pH | 7.0 | 7.0 | 7.0 | 7.0 | 7.0 | 7.0 | 7.4 | 7.4 | 7.4 | 7.4 |

### Reagents

Unless stated otherwise, reagents were from Sigma-Aldrich and were the highest available grade. 5 µM ionomycin free acid (Calbiochem) was used unless otherwise stated. Other reagents used were 5 µM FM4-64 (Life Technologies), 50-500 µM spermine (Sigma-Aldrich), and 100 µg/ml (0.01%) Trypan Blue (Sigma-Aldrich). Rhodamine-conjugated heptalysine (rhodamine-KKKKKKK-amide; K7) was prepared by Multiple Peptide Systems (NeoMPS, Inc.). Phosphatidyl-L-serine was from bovine brain (Sigma, P5660).

### Simulation of TMEM16F function and deep membrane compartments (Figs. 8 and 10)

Simulations of the working hypotheses, presented in Fig. 1, were written in Matlab. The fraction of TMEM16F channels that has no bound Ca^2+^ is ‘F_closed_’, the fraction that has no bound PS at regulatory sites is called ‘F_inactive_’, and the doubly inactivated fraction (F_closed_•F_inactive_) is not depicted in Fig. 1. The fraction of TMEM16F channels that is open (F_active_) can translocate cations via two-site pore, and can bind and translocate anionic phospholipids (PS) only when a cation is available at the relevant pore entrance.

In the initial state of all simulation, all PS (PS_tot_, 100 units) is assumed to be cytoplasmic:

1. PS_i_=100, and PS in the outer monolayer (PS_o_) is
2. PS_o_=PS_tot_-PS_i_ At any time, the fraction of active TMEM16F channels (F_active_, see Fig. 1B) is dependent on Ca^2+^ binding to a single TMEM16F site with a K_50_ of 20 µM and the binding of two PS molecules at regulatory sites. Total PS content of the membrane is always 100 PS units. The K_50_ for PS binding to activate TMEM16F is 120 PS units and the Hill slope is 2 (see Fig. 1B), implying that all TMEM16F channels are not activated by the normal PS content of the inner PM monolayer. Thus, the fraction of active TMEM16F channels (F_active_) is:
3. F_active_=Ca_i_/(Ca_i_+20µM) • PS_i_^2^/(PS_i_^2^+120^2^) The active state (F_active_) is then subdivided into conducting channels that have no PS bound within their open pores (E_o_), channels that have one PS bound within the pore on the cytoplasmic side (E_1_, and that have one PS bound within the pore on the extracellular side (E_2_). For simplicity, dual PS occupancy is not allowed in the simulations. The rates of PS translocation in TMEM16F (K_PSx_), PS binding by TMEM16F (K_PSon_) and dissociation from TMEM16F (K_PSoff_) are as follows, relevant to the total PS population of 100:
4. K_PSx_=20 s^−1^
5. K_PSon_=75 PS^−1^•s^−1^;
6. K_PSoff_=400 s^−1^ To simulate ion translocation within open TMEM16F channels, the E_0_ fraction of the F_active_ fraction of TMEM16F (see cation permeation model in Fig. 1), a two-site model is employed with all voltage dependence placed on ion translocation between inner and outer pore binding sites. The rates of ion binding (K_on_), dissociation (K_off_), and translocation between occupancy sites (K_x_) were assigned relative to a translocation rate (K_x_) of 1 s^−1^. K_on_ is 0.2 mM^−1^s^−1^ and K_off_ was 5 s^−1^. When changes of membrane potential (E_m_) were simulated, K_x_ was modified by a single, symmetrical barrier function,
7. K_em_ = e^(0.5•Em/26mV)^. With internal (X_i_) and external (X_o_) cation concentrations in mM, calculation of TMEM16F ion occupancy states and permeation rates proceed as follows (see Fig. 1B):
8. K_1_=X_o_•K_on_
9. K_2_=K_off_
10. K_3_ = K_off_
11. K_4_=X_i_•K_on_
12. K_5_= K_off_
13. K_6_=X_o_•K_on_
14. K_7_=X_i_•K_on_
15. K_8_= K_off_
16. K_9_=K_x_•K_em_
17. K_10_=K_x_/K_em_ Using a King-Altman procedure (Lam and Priest, 1972) to calculate occupancy of the four permeation states,
18. X_1_=K_2_•K_4_•(K_7_+K_6_)+K5•K_7_•(K_2_+K3)+K_10_•(K2+K_3_)•(K_7_+K_6_)
19. X_2_=K_1_•K_7_•(K_4_+K_5_)+K_4_•K_6_•(K_1_+K_8_)+(K_10_•K_1_+K_9_•K_4_)•(K_7_+K_6_)
20. X_3_=K_1_•K3•(K_7_+K_6_)+K8•K_6_•(K_2_+K3)+K_9_•(K_6_+K_7_)•(K_3_+K_2_)
21. X_4_=K_2_•K_8_•(K_4_+K5)+K3•K5•(K_1_+K_8_)+(K_9_•K_5_+K_10_•K8)•(K3+K_2_) With S=X_1_+X_2_+X_3_+X_4_, the fractional occupancies of the four states are:
22. F_1_=X_1_/S, F_2_=X_2_/S, F_3_=X_3_/S and F_4_=X_4_/S, where F_1_ is occupied on the cytoplasmic side by 1 cation, F_2_ by two cations, F_3_ by a cation on the extracellular side, and F_4_ is unoccupied. The (relative) permeation rate (R_net_) is then:
23. R_net_=F_3_•K_10_-F_1_•K_9_ Next, the active TMEM16F states with PS bound to the inner (E_1_ and outer (E_2_) pore binding sites are integrated over time. Assuming that PS can bind from the cytoplasmic side to all E_0_ states with cations occupying the cytoplasmic site (i.e. F1 + F2 states), assigning K_PSon_ (75 PS^−1^ • s^−1^) as the second order rate of PS binding, K_PSoff_ as the PS dissociation rate from E_1_ states (400 s^−1^), and K_PSx_ (20 s^−1^) as the PS translocation rate between PS sites in the pore,
24. dE_1_/dt = [PS_i_•E_0_•F_active_•K_PSon_•(F_1_+F_2_)-E_1_•K_PSoff_]+(E_2_-E_1_)•K_PSx_. To simulate the expectation that lipids in the outer monolayer are more ordered and less fluid, we assume that K_PSon_ is two-fold slower and that K_PSoff_ is two-fold faster on the extracellular side:
25. dE_2_/dt= [PS_o_•E_0_-F_active_•K_PSon_*(F_3_+F_2_)/2-E_2_*K_PSoff_•2]+(E_1_-E_2_)•K_PSx_. The fraction of active channels without a PS bound within the pore is calculated:
26. E_0_=1-E_1_-E_2_, PS changes in the cytoplasmic leaflet (PS_i_) are integrated,
27. dPS_i_/dt = E1•K_PSoff_ – PS_i_-E_0_ • K_PSon_ • F_act_ • (F_1_ + F_2_), PS in the extracellular leaflet is calculated,
28. PS_o_=PS_tot_PS_i_, the relative TMEM16F current is calculated,
29. I_TMEM16F_ = R_net_•F_active_•E_0_, and finally, the fraction of deep membrane compartments open to the extracellular space (F_open_) is calculated. We assume here that 4 PS molecules must be bound by the constricting membrane proteins and that the PS affinity is quite high. The K_50_, 35 PS units, is less than one-half of the total PS in the inner monolayer:
30. F_open_=35^4^/(PS_i_^4^+35^4^). Simulations were carried out in Matlab with accuracy checking to insure that simulation error was less than line-width in the results presented.

### Statistical analysis and replicate criteria

Statistical significance was evaluated with Students T-test. Results were rejected if a Shapiro-Wilk normality test failed. When outlier values were encountered, defined as experimental values that deviated more than two standard deviations from the mean, the values were removed from data sets if the number of experiments was greater than 6. Otherwise, the relevant experiments were repeated. All experiments described employed at least two batches of cells harvested on two different days.

### Supplemental Movie 1

TMEM16F-induced membrane expansion involves deep membrane compartments. As demonstrated in the accompanying article (Bricogne et. al., XXXX Figure 2), the impermeable styryl dye FM4-64 (3 µM) is employed to reversibly label the PM of Jurkat cells before and after membrane expansion. In the presence of physiological buffers (Solutions In_1_ and Out_1_) the Ca^2+^ ionophore ionomycin is used to elevate cytoplasmic Ca^2+^ and activate TMEM16F, causing a large increase in PM available for FM4-64 binding. Although FM4-64 fluorescence increases over-proportionally to C_m_, it is striking that large subsurface membrane compartments and invaginations become clearly visible and are labelled reversibly.

### Supplemental Movie 2

BHK-NCX cells were transiently transfected with VAMP4-pHlourin. After trypsinization, simultaneous whole-cell patch clamp and confocal imaging were used to monitor PM area, VAMP4-pHluorin fluorescence, TB (100 µg/ml) labelling of the PM, and TB quenching of VAMP4-pHluorin fluorescence. Solutions In_2_ and Out_2_ were employed, applying 2 mM extracellular Ca^2+^ to elevate cytoplasmic Ca^2+^ via NCX. Initially, VAMP4-pHluorin fluorescence (green) is not quenched by TB application, indicating that the fluorophore is inaccessible from the extracellular space. After Ca^2+^ elevation, a strong increase in VAMP4 quench indicates that VAMP4-pHluorin has become available to the extracellular medium. The incrementing VAMP4-fluorescence clearly colocalizes with deep, reversible TB labelling of the PM.

### Supplemental Movie 3

The cationic membrane probe K7r, which binds anionic phospholipids, reversibly labels deep PM compartments after cytoplasmic Ca^2+^ elevations. Similar to experiments in Fig. 11, BHK-NCX cells were patch clamped and C_m_ was monitored. Simultaneously, confocal imaging was carried out to detect PS exposure via K7r fluorescence. Phospholipid scrambling was induced by using Solution Out_3_ with 140 Cs and Solution In_5_ with 120 TEA, set to ~5 µM free Ca^2+^. According to the working hypothesis, phospholipid scrambling takes place by the forward electrogenic scrambling mode. K7r (3 µM) was periodically applied and washed off via extracellular solution switches. Membrane expansion occurs in parallel with extracellular exposure of anionic phospholipid, as detected by K7r fluorescence. Similar to staining by the other membrane probes, K7r fluorescence reveals deeply invaginating membrane compartments.

